# Bivariate GWAS scan identifies six novel loci associated with lipid levels and coronary artery disease

**DOI:** 10.1101/319848

**Authors:** Katherine M. Siewert, Benjamin F. Voight

## Abstract

**Background:** Plasma lipid levels are heritable and genetically associated with risk of coronary artery disease (CAD). However, genome-wide association studies (GWAS) routinely analyze these traits independently of one another. Joint GWAS for two related phenotypes can lead to a higher-powered analysis to detect variants contributing to both traits.

**Methods and Results:** We performed a bivariate GWAS to discover novel loci associated with heart disease, using a CAD Meta-Analysis (122,733 cases and 424,528 controls), and lipid traits, using data from the Global Lipid Genetics Consortium (188,577 subjects). We identified six previously unreported loci at genome-wide significance (P < 5 × 10^−8^), three which were associated with Triglycerides and CAD, two which were associated with LDL cholesterol and CAD, and one associated with Total Cholesterol and CAD. At several of our loci, the GWAS signals jointly localize with genetic variants associated with expression level changes for one or more neighboring genes, indicating that these loci may be affecting disease risk through regulatory activity.

**Conclusions:** We discovered six novel variants individually associated with both lipids and coronary artery disease.

## Introduction

Genome-wide association studies (GWAS) for coronary artery disease (CAD) have identified numerous susceptibility loci associated with the disease.^1–3^ These loci account for only a fraction of the total heritability for CAD, indicating that additional loci remain to be discovered.^4^ For many CAD-associated regions, an obvious mechanistic connection to heart disease is not clear. To identify novel mechanistic connections, a recent trend is to utilize *data integration*, whereby associations for a phenotypic endpoint (like CAD) are jointly statistically analyzed with additional data types to better understand the underlying phenotypic association. This raises two questions, namely: (i) which datasets should be used, and (ii) what statistical procedures should be employed for data integration?

The community is poised to make substantial progress in the understanding of CAD through integration of diverse, publicly available genome-wide datasets. This is due in part to a basic understanding of heritable risk factors that contribute risk to CAD.^5^ Specifically, it is known that low-density lipoprotein cholesterol (LDL-C) is a causal risk factor for CAD, and emerging evidence supports elevated triglycerides as an additional causal risk factor.^6–10^ Because serum lipid levels are heritable, large-scale association scans for these traits are available, and thus can be readily integrated with CAD association data.^11,12^ In addition, large-scale studies have now emerged which map genetic variation associated with changes in gene expression *(i.e*., eQTLs) across major human organs and tissues.^13^ These data provide an opportunity to link a CAD association signal to a putative causal gene and potentially a tissue of action.

To address the second question, several new statistical approaches have emerged. Multivariate studies that combine association data from multiple traits can help interpret the underlying associations at existing loci and have higher power for novel locus discovery than univariate studies.^14,15^ We and others have demonstrated the utility of such approaches through their use in identifying novel loci affecting CAD and type 2 diabetes,^15^ or loci affecting fasting insulin, triglyceride levels and HDL cholesterol levels.^16^ A key advantage of this approach is the potential for direct mechanistic insight into how a locus may affect the phenotypic endpoint of interest. For example, a locus associated with both cholesterol and heart disease might suggest a mechanism of action and therapeutic hypothesis to lower disease risk. However, coincidental associations of two traits at a genomic locus may not reflect a unified etiology. Thus, new methods have been developed which quantify the evidence that an overlapping association pattern across two datasets reflects a single underlying association *(i.e*., statistical colocalization), instead of coincidental overlap in which the associations are due to different causal variants.^17^

Here, we report a bivariate GWAS scan to discover novel loci associated with CAD and four lipid-related traits: LDL cholesterol (LDL-C), HDL cholesterol (HDL-C), Total Cholesterol (TC) and Triglycerides (TG) levels. To focus on a set of loci that potentially point to a shared causal variant and cognate gene of interest across traits, we applied statistical colocalization for the lipid and CAD association signals with each other and with gene expression datasets. We report six novel genome-wide significant loci where the GWAS association signals statistically colocalize: three affecting CAD and TG levels, two affecting CAD and LDL-C levels and one affecting CAD and TC. Of these loci, four signals also colocalized with eQTL signals from GTEx,^13^ indicating gene expression may be influencing disease risk at these loci.

## Results and Discussion

### Overview of Bivariate Scan

Using a bivariate GWAS scan approach, we identified six novel loci associated with lipid levels and heart disease. As input into this scan, we used data from a meta-analysis of coronary artery disease, which combined results from the CARDIOGRAM+C4D consortium (88,192 CAD cases and 162,544 controls)^2^ and the UK BioBank (34,541 cases and 261,984 controls)^3^ and GWAS results for each of four lipid traits: HDL-C, LDL-C, Total Cholesterol, and Triglycerides, from the Global Lipid Genetics Consortium (188,577 individuals).^12^ Our trait covariances and bivariate p-values appeared calibrated (**Supplementary Table 1**, **Supplementary Figure 1**) and had a strong correlation with single-trait P-values, as would be expected (**Supplementary Figures 2 and 3**). In addition, the top hits from our scan included established lipid and CAD associated loci, including *PCSK9* (significant for CAD paired with LDL-C and TC), *APOE* (CAD paired with all four lipids traits), and *LPL* (CAD paired with HDL-C and TG).

We performed several filtering steps to identify variants with non-trivial associations (P < 5 × 10^−8^) not yet established from individual trait scans. We first removed loci from our set with prior reported associations with either CAD or the lipid trait under consideration.^1,11,18–20^ Next, we narrowed our set of variants to those that were nominally associated with both traits from single-trait association data (P < 5 × 10^−3^ for the lipid trait and CAD). To focus on sites where we could hypothesize that the same causal variant contributes to both the lipid and CAD signals, we filtered out variants where the patterns of association for both traits did not statistically overlap (COLOC PP4/(PP3+PP4) < 0.5, **Methods, Supplementary Table 2**). After these filters, we observed two loci associated with LDL-C and CAD, three with CAD and Triglycerides and one with CAD and TC (**Table 1**). Sentinel SNPs at these loci were not strongly linked to a nonsynonymous variant (**Methods**), consistent with a gene regulatory mechanistic hypothesis to explain phenotypic variability.^21^ We subsequently focus on associations with eQTLs or relevant prior phenotypic connections. One additional lead SNP, rs7033354, is associated with LDL-C and CAD (P = 1.16 × 10^−8^, **Table 1**, **Supplementary Figure 4**).^22^

**Table 1:** Significant loci by bivariate scan with colocalization of the CAD and lipid trait GWAS signals

| Variant | chr | Position (Mb) | Lipid Trait | EA/OA | EAF | Bivariate P | Lipid Beta | Lipid s.e. | Lipid P | CAD OR | CAD 95% CI | CAD P |
| --- | --- | --- | --- | --- | --- | --- | --- | --- | --- | --- | --- | --- |
| rs895953 | 12 | 122.2 | TG<sup>†</sup> | T/G | 0.81 | 2.4x10<sup>-9</sup> | 0.022 | 0.0043 | 8.6x10<sup>-8</sup> | 0.98 | 0.97-0.99 | 9.1x10<sup>-4</sup> |
| rs3741782 | 12 | 109.0 | LDL | A/G | 0.69 | 3.2x10<sup>-9</sup> | -0.020 | 0.0039 | 6.3x10<sup>-7</sup> | 0.97 | 0.96-0.99 | 3.3x10<sup>-6</sup> |
| rs7033354 | 9 | 16.9 | LDL | T/C | 0.63 | 1.2x10<sup>-8</sup> | -0.019 | 0.0038 | 1.4x10<sup>-6</sup> | 0.98 | 0.97-0.99 | 7.4x10<sup>-6</sup> |
| rs12423664 | 12 | 133.0 | TG | A/G | 0.13 | 1.2x10<sup>-8</sup> | 0.026 | 0.005 | 8.2x10<sup>-8</sup> | 1.03 | 1.01-1.05 | 5.2x10<sup>-4</sup> |
| rs2908290 | 7 | 44.2 | TG | A/G | 0.38 | 1.6x10<sup>-8</sup> | 0.017 | 0.0034 | 1.5x10<sup>-6</sup> | 1.02 | 1.01-1.03 | 1.4x10<sup>-5</sup> |
| rs1378942 | 15 | 75.0 | TC | A/C | 0.62 | 2.2x10<sup>-8</sup> | 0.012 | 0.0037 | 9.1x10<sup>-6</sup> | 0.98 | 0.97-0.99 | 1.8x10<sup>-5</sup> |

\*EA: Effect Allele, OA: Other Allele, OR: Odds Ratio, CI: Confidence Interval, EAF: Effect Allele. Position is aligned to hg19. Frequency (EUR in 1000 Genomes).
<sup>†</sup> In addition to the TG association, this SNP is nominally associated with HDL (Bivariate P = 8.78 x 10<sup>-8</sup>)

### The monogenic diabetes gene glucokinase is associated with TG and CAD

Our first variant, rs2908290, is associated with TG and CAD (Bivariate P = 1.6 × 10^−8^) and is in an intron of the GCK (glucokinase) gene (**Table 1**, **Figure 1)**. Although this SNP is not an eQTL in any assayed tissue, prior evidence supports GCK as the causal gene at this locus. Glucokinase phosphorylates glucose as the first step of glucose metabolism and is an established gene for maturity-onset diabetes of the young (MODY), a monogenic disorder causing early onset diabetes.^23^ It has been robustly associated with metabolic phenotypes in GWAS, including fasting glucose, glycated hemoglobin and type 2 diabetes (T2D).^15,19,24,25^ A mutation in the promoter of *GCK* was previously found to be associated with CAD in a candidate gene analysis,^26^ however, their variant is in very low linkage with our lead variant (European LD r^2^=0.05).

**Figure 1:**
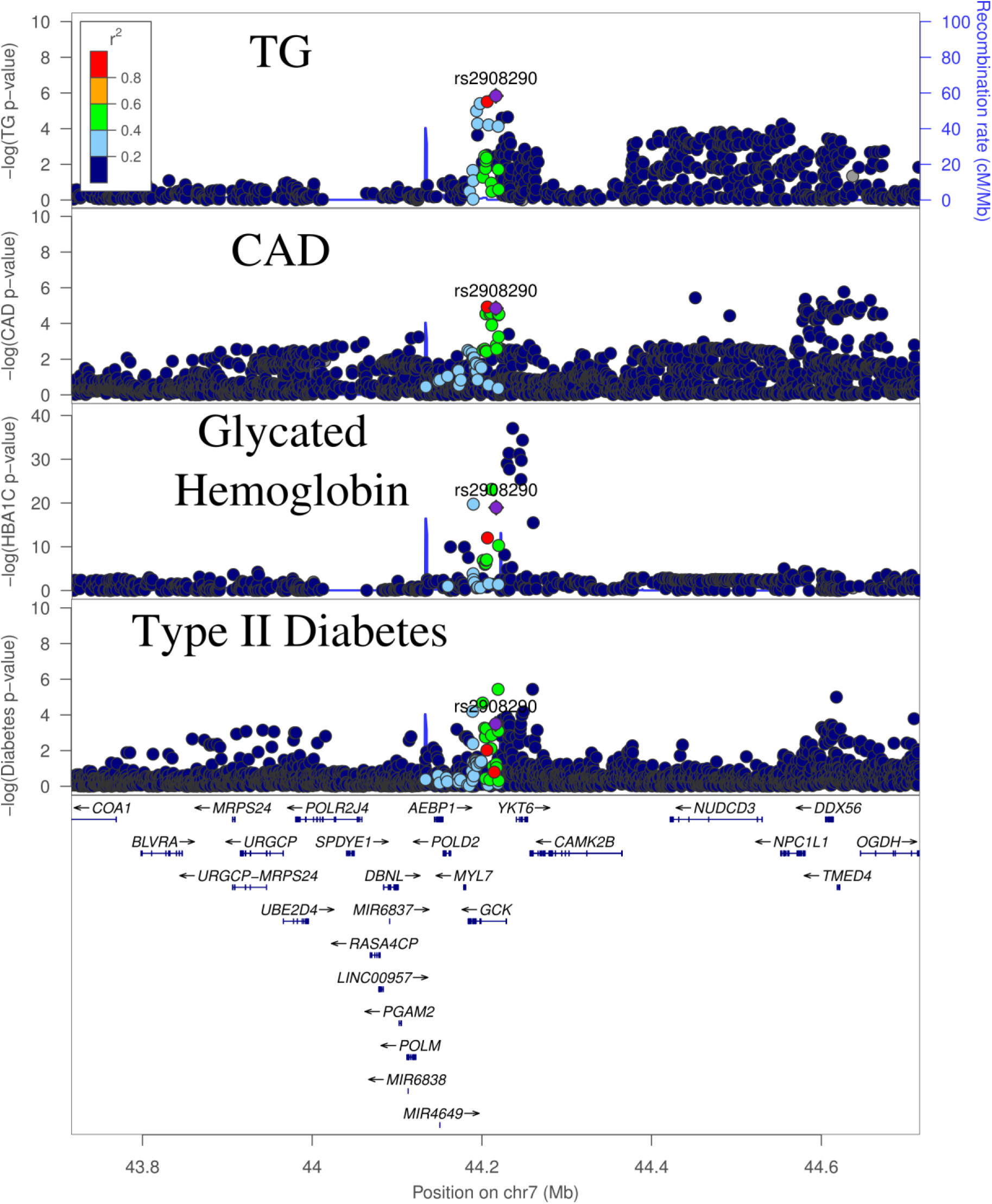
GWAS signals for Triglycerides (TG), CAD and T2D at the GCK locus.^15,25^

The association signals at this locus are complex and suggest the possibility of several causal variants. Bien et al. found evidence of two independent associations at this locus for fasting glucose, with rs2908290 as the lead SNP of the second association.^27^ In line with this, we found evidence of three independent genome-wide significant causal variants for glycated hemoglobin, a marker of long-term elevated glucose, within 100kb of our lead SNP using approximate conditional analysis (**Methods**).^25^ However, none of the three lead variants are strongly linked with rs2908290, indicating the possibility of many different casual variants for various metabolic traits at this locus. To examine if the TG and CAD signal at rs2908290 statistically overlap, we performed a colocalization analysis on a window 100kb to either side of this SNP, to exclude a nominally significant SNP several hundred kb upstream of *GCK*. The TG and CAD GWAS showed strong evidence of statistical overlap (COLOC PP4 = 0.87). Long-term overexpression of GCK has been shown to cause both hyperinsulinemia and hypertriglyceridemia in mice^28^, which is broadly consistent with the pattern of genetic associations observed at this site (**Supplementary Table 3**).

As it is known that *GCKR* (glucokinase regulator) inhibits the activity of *GCK*, we next investigated the CAD association at the common variant in this gene, Pro446Leu. Previous functional studies have shown that this variant diminishes the capability of GCKR to inhibit GCK, which results in increased glycolic flux and liver metabolites that may promote *de novo* lipogenesis^29^, providing a mechanism to explain the observed human genetic associations of decreased fasting glucose and T2D risk but increased triglyceride levels^12,15,24,29,30^. In addition to these established associations, we observed a modest association of higher CAD for carriers of the triglyceride-increasing allele (OR = 1.018, P = 1.9 × 10^−3^, **Supplementary Table 3**). This observed effect is consistent with the predicted combined effect of elevated triglycerides and lower fasting glucose levels on CAD risk (Predicted OR = 1.022, **Methods**). These data cumulatively suggest that genetic disruption of *GCK* influences diabetes and cardiovascular disease risk through known, causal intermediates.

### Locus associated with TG and HDL-C may regulate genes affecting tyrosine catabolism or platelet traits

The second locus is associated in our bivariate scan with both TG and CAD (Bivariate P = 2.35 × 10^−9^), with HDL-C and CAD (Bivariate P = 8.78 × 10^−8^, **Figure 2**), and is also an eQTL for several nearby genes. The lead variant, rs895953, is in an intron of the *SET1B* gene and, to our knowledge, has no prior established GWAS associations for related traits. Our sentinel SNP is an eQTL for three genes in the region in multiple tissues (**Supplementary Table 4)** including *RHOF* (Ras Homolog Family Member F, Filopodia Associated) and *HPD* (4-Hydroxyphenylpyruvate Dioxygenase). In contrast to epidemiological expectation, the major allele of this variant is associated with an increase in Triglycerides, but a reduced risk of CAD.

**Figure 2.**
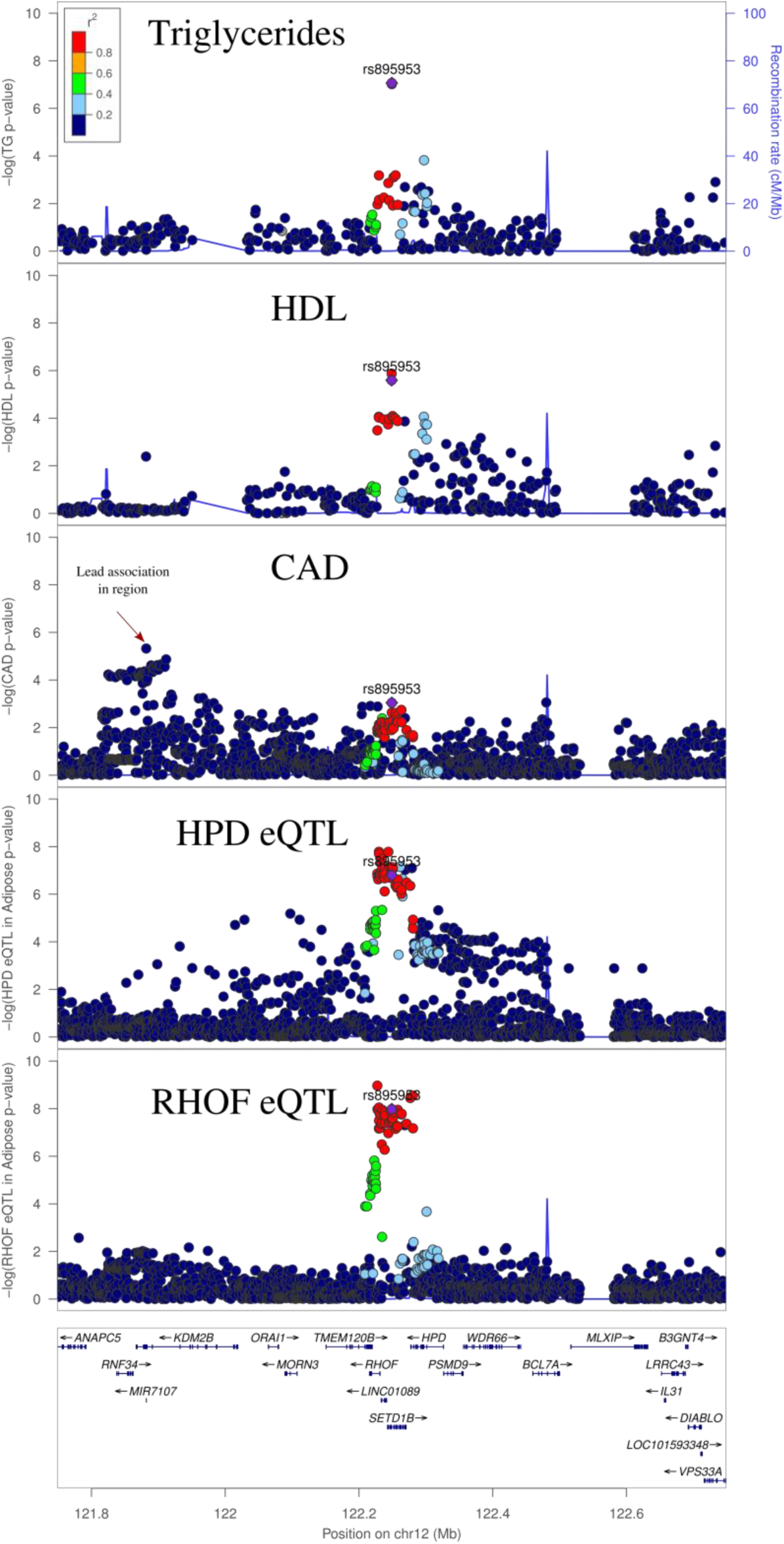
GWAS and eQTL associations at the *SET1B* locus. Triglyceride, HDL-C and CAD signals are the GWAS signal corresponding to the respective traits. eQTL associations are for expression of the gene in Adipose Subcutaneous.

There is strong evidence of colocalization between *RHOF* and *HPD* expression and the TG signal (**Supplementary Table 4)**. The CAD colocalization was weaker due to an unlinked CAD signal nearby (European LD r^2^=0.0, **Figure 2**, 3^rd^ panel denoted by the arrow) which would violate the assumptions of the approach used to quantify colocalization. To confirm that the CAD signal overlapping the lipids and eQTL signals was not due to this secondary signal, we performed an approximate conditional analysis (**Methods**), conditioning on the top SNP from the nearby peak (rs10849885). The association strength for our sentinel SNP improved slightly (P_unconditional_ = 9.1 × 10^−4^ to P_conditional_ = 3.6 × 10^−4^) upon conditioning, consistent with the lack of linkage disequilibrium between our sentinel SNP and the secondary CAD signal. Furthermore, the probability of colocalization between the eQTLs for *RHOF, HPD*, and each GWAS signal, as well as between the lipid and CAD GWAS signals themselves, also increased when we performed a colocalization analysis using a 200kb window in this region, which excludes the secondary CAD signal (**Supplementary Table 2, Supplementary Table 4**). These results are consistent with the hypothesis that a single causal variant underlies the GWAS signals and the expression association for *RHOF* and *HPD* at this locus.

Review of the literature suggests that one or both genes are plausible candidate genes for the CAD and lipid associations. *HPD* is involved in the catabolism of tyrosine, and has been associated with 2-Hydroxyisobutyrate levels, though this previous variant is not strongly linked with ours (European LD r^2^=0.03 between our lead SNP and that from Suhre and Raffler et al.),^31,32^. 2-Hydroxyisobutyrate concentration is a known biomarker for insulin resistance and adiposity.^33,34^ However, literature may also support the hypothesis that *RHOF* as causal. This gene lies in an GWAS association for platelet count, platelet volume, and reticulocyte fraction of red cells.^22^ Platelet biology has been implicated in lipid metabolism and atherosclerosis, suggesting rs895953 could be affecting CAD risk through platelet-related pathways.^35,36^

### Locus associated with LDL-C and CAD may affect actin remodeling

A locus tagged by the sentinel SNP rs3741782 was associated with LDL-C and CAD (Bivariate P = 3.2 × 10^−9^) and fell within an intron of the *CORO1C* gene *(i.e*., Coronin, Actin Binding Protein 1C, **Figure 3**). This SNP is an eQTL for *CORO1C* in visceral adipose tissue and in the adrenal gland, and for *SSH1 (i.e*., Slingshot Protein Phosphatase 1) in whole blood, with strong evidence of colocalization of theses eQTL peaks with the CAD and LDL-C GWAS signals (**Supplementary Table 4**). Both *SSH1* and *CORO1C* regulate actin reorganization.^37,38^ It has been demonstrated that uptake and degradation of lipoproteins by macrophages requires an actin cytoskeleton,^39^ and oxidization of the associated lipids in macrophages can result in foam cells, which contribute to atherosclerosis.^40^ However, to our knowledge, it is unknown whether *CORO1C* or *SSH1* participate in this mechanism.

**Figure 3:**
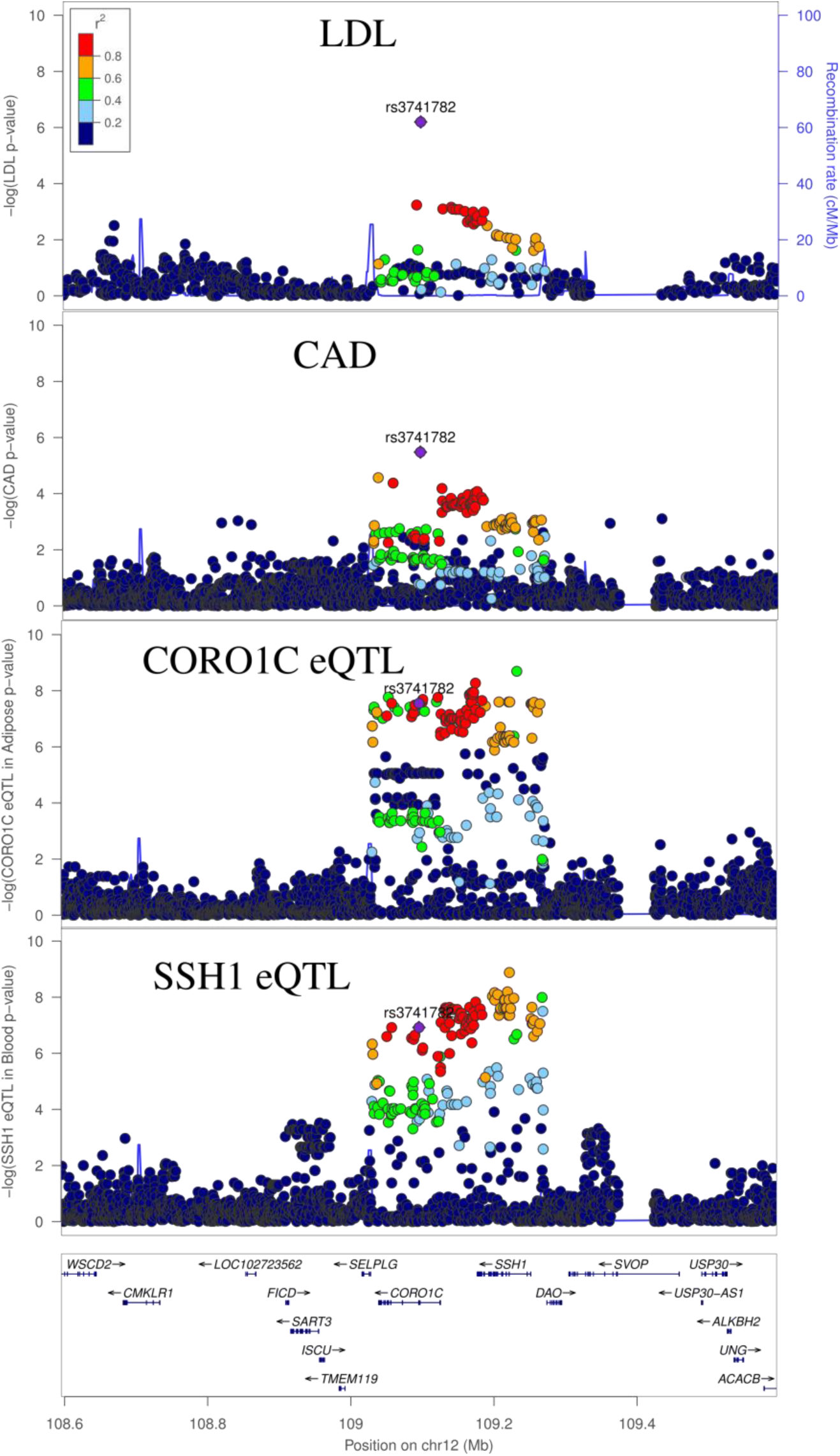
GWAS and eQTL Associations at the CORO1C locus. LDL-C and CAD signals are the GWAS signal corresponding to the respective traits. CORO1C eQTL signal is for expression in adipose, and SSH1 is for expression in whole blood.

**Figure 4:**
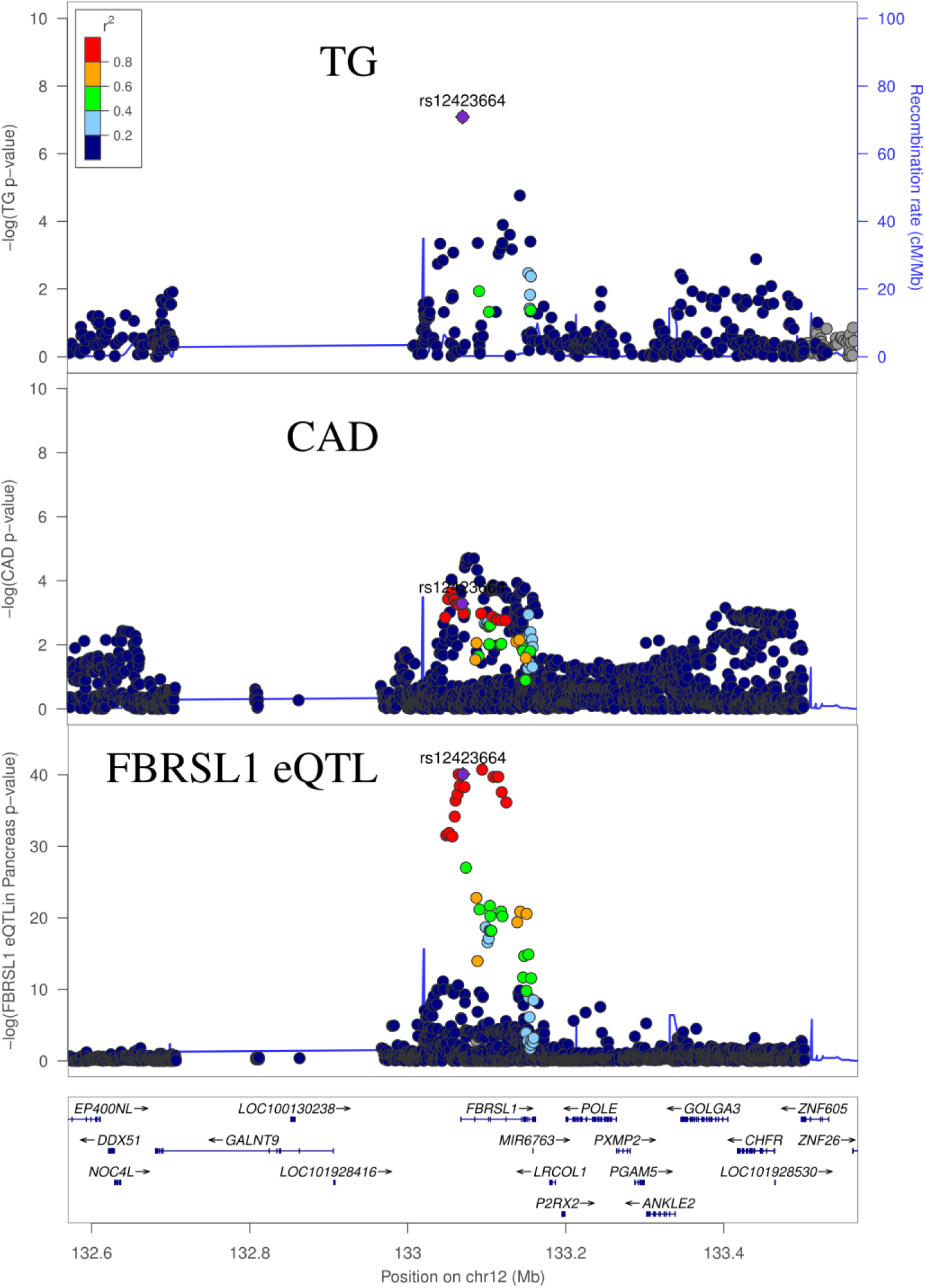
GWAS and eQTL Associations at the FBRSL1 locus. Triglyceride (TG) and CAD signals are the GWAS signal corresponding to the respective traits. FBRSL1 eQTL is signal is for expression of the gene in pancreas.

### Locus associated with TG and CAD may regulate expression of *FBRSL1*

A fourth locus, tagged by sentinel SNP rs12423664 (Bivariate P = 1.2 × 10^−8^), is located in an intron of *FBRSL1* (i.e., Fibrosin Like 1). Evidence has shown that the protein encoded by this gene may be a member of the polycomb repressive complex 1 *(PRC1)*, which regulates gene expression via epigenetic markers.^41,42^ Our lead SNP is an eQTL for this gene in both pancreas and whole blood. Our sentinel variant is partially linked to a previous GWAS association with red blood cell count (European LD r^2^=0.52 between rs34390795 and our sentinel SNP).^22^

Available evidence suggests that the eQTL association may underlie the lipid and CAD association. Colocalization analysis supports a shared underlying causal variant between the lipid, CAD signal, and the eQTL (**Supplementary Table 1, Supplementary Table 4**). That being said, the underlying lipid GWAS was not as density genotyped as the CAD and eQTL scans (**Figure 3**). Indeed, the lead GWAS variant for CAD (Eueopean LD r^2^=0.08 between rs4883525 and rs12423664), along with several other top CAD variants were not assayed in the lipid GWAS data. We expect larger-scale imputation studies for the lipids scan should enable more well-powered colocalization analysis at this locus.

The Fibrosin-Like 1 protein has been found to have lower expression in human hearts with dilated cardiomyopathy, relative to control hearts.^43^ In the eQTL signal colocalizing with our bivariate peak, the alternative allele (A) of our lead SNP is associated with lower expression of this gene, though this association was observed in pancreas, not heart.^13^ This same allele is associated with increased lipid levels and an elevated risk of heart disease. These results indicate that Fibrosin Like 1 may play a role in healthy heart processes, and thus reduced expression of this gene may increase risk of heart disease. However, the diverse biological processes that *PRC1* may influence make it difficult to interpret how this gene may be affecting triglycerides and heart disease.

### Variant affecting blood pressure, CAD and Total cholesterol

Our final variant, rs1378942, is associated in our bivariate scan with CAD and total cholesterol (Bivariate P = 2.22 × 10^−8^), and also nominally associates with LDL-C (**Figure 5**). The lipid traits strongly colocalize with each other and moderately colocalize with the CAD locus (**Supplementary Table 4)**. In addition, this index SNP has an established association with systolic blood pressure (P = 6 × 10^−23^),^44^ with moderate to strong evidence of colocalization between both lipid traits, CAD and SBP (**Supplementary Table 5**). The ‘A’ allele of this SNP increases LDL and TC while decreasing SBP and CAD, pointing to SBP as the directionally consistent causal phenotype for the CAD association.

**Figure 5:**
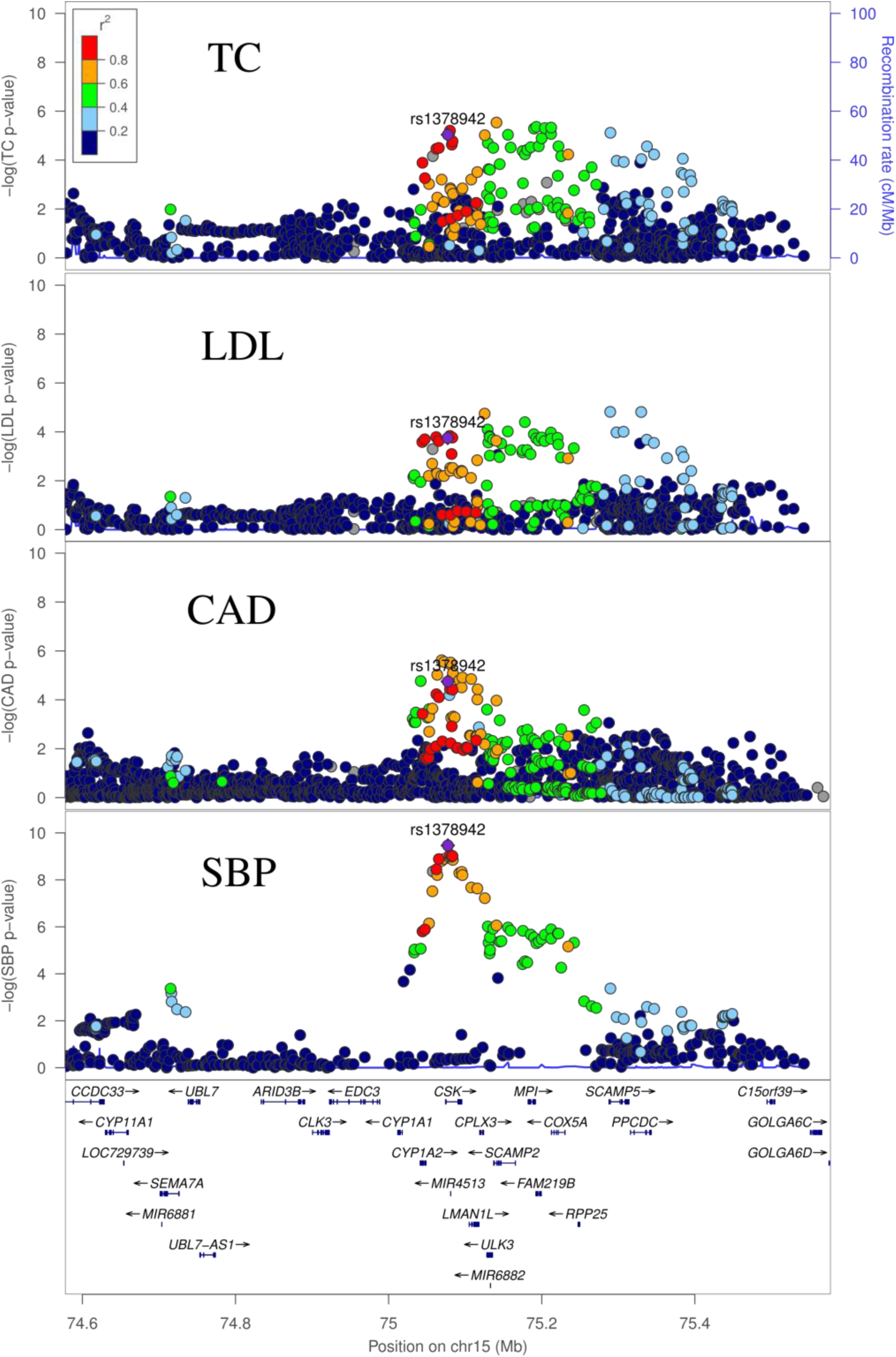
GWAS Associations at the CSK locus. Total Cholesterol (TC), LDL-C and SBP (systolic blood pressure) and CAD signals are the GWAS signal corresponding to the respective traits.

rs1378942 is an eQTL for *CSK* (C-terminal Src Kinase), in both subcutaneous adipose and the left ventricle of the heart, and *CYP1A1* (Cytochrome P450 Family 1 Subfamily A Member 1) in Subcutaneous Adipose and Muscle Skeletal, among several other genes.^13^ The eQTL signals for these two genes have moderate to strong evidence of colocalization with the CAD and TC GWAS peaks, although visual inspection shows that there may be more than one causal eQTL variant for the *CSK* eQTL association (**Supplementary Figure 5, Supplementary Table 4**). siRNA knockdown of *CSK* was found to increase blood pressure, suggesting it is as a possible causal gene for SBP at this locus.^45^ However, *CYP1A1* is another possible causal gene. Although the substrate of the gene is not firmly established, members of the cytochrome p450 family are known to affect cardiovascular disease risk through effects on a diverse set of processes, including lipid metabolism and vasoconstriction.^46^

## Conclusion

Using a bivariate approach, we identified six novel variants associated with both CAD and one or more lipid traits, demonstrating the power of bivariate GWAS approaches to discover novel loci influencing phenotypes of interest, with potential candidate genes identified via integration with gene expression data. These loci are of particular interest, because they provide human genetic association evidence of altering CAD risk through lipid levels. However, other scenarios are also possible: namely, a variant may not affect CAD directly through lipid levels, and could instead influence other processes by which lipid and CAD risk levels are altered independently of one another.

Our results speak to the complexity of interpreting functional mechanisms of associated variants. Three of our loci suggest the possibility of several different regulatory targets, indicating these loci may not affect CAD and lipids through a single regulatory mechanism. Instead, these loci may harbor regulatory variants for several genes, which independently affect lipid levels and therefore CAD. Alternatively, as shown in the case of rs3741782, these regulatory targets may be involved in the same biological pathway. Furthermore, conditional analyses reveal that several of the GWAS and eQTL signals may harbor more than one causal variant, as has been previously demonstrated for several metabolic-related loci.^47,48^

The combination of lipid and heart disease associations and their colocalization with eQTLs for genes involved with cardiometabolic process supports the potential therapeutic potential of the loci found in our bivariate scan. Further functional experimental work will be required to verify and elucidate these candidates’ association with lipid and CAD pathways.

## Methods

### GWAS Data Collection

Our CAD GWAS data was from a meta-analysis that was performed using the UK BioBank and several prior CAD GWAS studies.^3^ Lipid GWAS data was obtained from the Global Lipids Genetics consortium.^12^ The joint analysis of Metabochip and GWAS data was used for all four lipid traits.

To generate z-scores for the lipid traits with genomic control correction, we used the following expression:

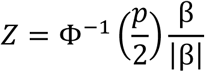

where Φ^−1^ is the inverse-cumulative distribution function of the normal distribution, p is the genomically-controlled p-value and *β* is the effect size of the single-trait association, as was done in Zhao et al..^15^

### Bivariate Scan

To perform the bivariate scan, alleles were first harmonized using the harmonise_data function in MRbase (https://github.com/MRCIEU/TwoSampleMR). A set of LD-independent variants was obtained using plink with the command --indep-pairwise 1000 5 0.2, ^49^ using 1000 Genomes phase 3 data from Europeans for linkage information. This set of independent variants was used to estimate the means and covariance matrix of the bivariate normal distribution of the z-scores. These parameters were then used to calculate a p-value for each SNP contained in both GWAS datasets using a chi-squared test with two degrees of freedom (code available at https://github.com/WWinstonZ/bivariate scan).

Clusters of significantly associated variants were produced using the --clump-r2 0.2 command in R. To focus on novel loci, we filtered out variants that had a P < 5 × 10^−8^ for either trait in our GWAS datasets, as these represented known loci. We additionally filtered out variants with a P > 5 × 10^−3^ in one of the two traits to focus on variants with convincing evidence of association with both traits. We further removed GWAS variants that were within 1MB and r^2^ > 0.2 of a significant loci associated with CAD or the relevant lipid trait in the GWAS catalog (downloaded 2/20/18),^19^ Nelson et al.,^1^ Klarin et al.,^20^ Lu et al.^18^ or Liu et al.^11^ We used the annovar package to search for functional mutations in r^2^ >= 0.8 of each lead SNP.^50^

### GTEx Data Collection

eQTL data from the GTEx consortium version 7 was downloaded for the following tissues: Adipose Subcutaneous, Adipose Visceral Omentum, Artery Aorta, Artery Coronary, Artery Tibial, Heart Atrial Appendage, Heart Left Ventricle, Liver, Muscle Skeletal, Pancreas, Thyroid and Whole Blood.^13^ We focused on this subset of tissues, as (i) the most plausible for our collection of phenotypes and disease endpoints, and (ii) eQTL discovery power, based on the sample size.

### Locus Characterization

We performed colocalization analysis of the GWAS using the coloc.abf() function from the coloc package (https://github.com/chr1swallace/coloc).^17^ A threshold of PP4/(PP3+PP4) greater than 0.5 was used for colocalization, with default region size of 500kb on either side of the SNP of interest.

We also conducted a colocalization analysis between eQTL signals and our top bivariate GWAS hits using coloc. We performed coloc analysis between both the lipid GWAS and the eQTL signal, and the CAD GWAS and the eQTL signal. A threshold of PP4/(PP3+PP4) ≥ 0.8 was used for “strong” evidence of colocalization and ≥ 0.5 was used for “moderate” evidence. We performed the coloc analysis on any significant eQTL signal in GTEx at our bivariate SNP of interest in any of the aforementioned tissues.

Linkage disequilibrium for our analyses other than those using plink was calculated with LDlink (https://analysistools.nci.nih.gov/LDlink/) using EUR populations from the 1000 Genomes Project.^51^ Locus plots were generated using LocusZoom also using EUR populations.^52^

We used the COJO method,^53^ implemented in GCTA,^54^ to perform our conditional analyses. In each case, we conditioned the lead GWAS SNP in our bivariate scan on the lead SNP of the nearby GWAS signal from which we were testing independence. To estimate the number of independent effects for the GCK locus, the stepwise model selection method in COJO with default parameters was used.

### Predicted effect of GCKR on CAD risk

We calculated the expected change in CAD risk due to the T2D and TG associations of rs1260326 at the GCKR locus. The odds ratio of CAD associated with 1 standard deviation change in triglycerides has been estimated as 1.28,^55^ corresponding to a β of ln(1.28)=0.247. The corresponding change in CAD risk accompanying the TG β of this SNP of −0.11 (**Supplementary Table 3**) is then −0.11 * 0.247 = –0.027. The effect of a 1 standard deviation change in T2D on CAD risk is estimated to be 1.11,^56^ corresponding to a β of ln(1.11) = 0.104. The 0.049 standard deviation change in T2D risk associated with rs1260326 is then expected to cause a CAD β of 0.104*0.049=0.0051. Assuming an additive model, the predicted effect of rs1260326 on CAD is then −0.027 + 0.0051 = −0.022, which is close to the observed effect of −0.018, corresponding to an odds ratio of 0.98.

### Funding Sources

This work was supported by a grant from the National Institutes of Health NIDDK R01DK101478 to B.F.V.

